# An inflammation-associated five-gene expression signature stratifies survival and immune states in lung adenocarcinoma: an integrative public-cohort analysis

**DOI:** 10.64898/2026.08.21.746098

**Authors:** Xingchen Zhou, Zhen Le, Pengxia Song, Qianru Xu, Ming Chen, Xiaobo Liu, Mengnan Cao, Sijie Zhan, Yubin Liu, Lin Zhang

## Abstract

**Background:** Inflammation and the tumor immune microenvironment contribute to lung adenocarcinoma (LUAD) progression, but the relationship among inflammation-linked transcriptional heterogeneity, patient survival, and immune-state variation remains incompletely defined.

**Objective:** We aimed to identify inflammation-associated LUAD subtypes, derive a parsimonious survival-stratification signature, and characterize its immune and pathway context across public transcriptomic cohorts.

**Methods:** Expression profiles and clinical data were obtained from TCGA-LUAD, GTEx normal lung, and GEO datasets GSE11969, GSE30219, GSE31210, and GSE40791. A curated set of 596 inflammation-related genes was used for consensus clustering. Differential-expression analysis, functional enrichment, univariate Cox regression, and LASSO-Cox modeling were integrated to construct a gene-expression risk score. The prognostic dataset comprised 730 cases and was randomly divided into training (n=502) and internal-validation (n=228) sets; 85 GSE30219 cases formed an external-validation cohort. Immune-cell enrichment, gene set enrichment analysis (GSEA), gene set variation analysis (GSVA), and pan-cancer analyses were used for biological contextualization.

**Results:** The LUAD-versus-control comparison identified 1,305 differentially expressed genes, including 498 upregulated and 807 downregulated genes. Consensus clustering resolved two inflammation-associated subtypes and 67 subtype-associated genes, of which 64 were higher and 3 were lower in Cluster 1 relative to Cluster 2. Thirty-three genes overlapped between the tumor-control and subtype contrasts. LASSO-Cox regression selected *CHRDL1, FDCSP, CXCL13, CYP4B1*, and *S100P*. The 1-, 3-, and 5-year areas under the time-dependent receiver operating characteristic curve were 0.6625, 0.6581, and 0.6658 in the training set; 0.7422, 0.6537, and 0.6761 in internal validation; and 0.6560, 0.6387, and 0.6753 in external validation. Risk groups differed across multiple T-cell, B-cell, natural-killer-cell, myeloid, dendritic-cell, macrophage, and granulocyte signatures. Positive GSEA signals included cell cycle (normalized enrichment score [NES]=2.67; adjusted P=1.42×10^−8^), DNA replication (NES=2.52; adjusted P=2.52×10^−7^), and mismatch repair (NES=2.20; adjusted P=1.77×10^−4^).

**Conclusions:** The five-gene expression score separated LUAD survival groups and captured coordinated proliferative and immune transcriptional states. Its moderate discrimination supports further biological and clinical validation rather than immediate clinical application.

**Brief Summary:** Public LUAD transcriptomes identified two inflammation-associated subtypes and a five-gene score comprising *CHRDL1, FDCSP, CXCL13, CYP4B1*, and *S100P* that separated survival groups and marked distinct proliferative and immune expression programs.

## Introduction

Lung cancer remains a leading cause of cancer mortality worldwide, and lung adenocarcinoma (LUAD) is the most common histologic subtype of non-small-cell lung cancer [1,2]. Large-scale genomic and transcriptomic studies have established that LUAD comprises molecularly diverse tumors rather than a single biological entity [3]. This heterogeneity influences disease progression, therapeutic sensitivity, and survival, creating a need for molecular classifications that complement clinicopathologic staging.

Inflammation is an enabling feature of cancer and a major determinant of host-tumor interaction. Chronic inflammatory signaling can support malignant-cell proliferation, survival, angiogenesis, extracellular-matrix remodeling, invasion, and immune suppression [4-6]. At the same time, the composition and organization of immune cells within tumors can be associated with clinical outcome and treatment response [7-9]. Stromal and extracellular-matrix programs further modulate tumor-cell behavior, metastatic competence, and immune access [10,11]. Bulk transcriptomic profiles therefore capture a composite state contributed by malignant, stromal, and immune compartments.

A large number of expression-based prognostic signatures have been proposed for LUAD, but many remain difficult to interpret or reproduce because feature selection is detached from a defined biological hypothesis, platform-specific preprocessing is incompletely reported, and clinical validation is limited. An inflammation-focused framework may improve interpretability by connecting tumor dysregulation with intratumoral immune heterogeneity, but it is important to distinguish statistical association from cellular mechanism and clinical utility.

We integrated public TCGA, GTEx, and GEO transcriptomic resources to characterize LUAD-versus-control expression differences, define inflammation-associated tumor subtypes, and identify genes shared by tumor dysregulation and subtype heterogeneity. Survival-associated genes were reduced by univariate Cox and LASSO-Cox regression to a five-gene expression score. The score was evaluated in training, internal-validation, and external-validation cohorts, followed by immune-cell enrichment, GSEA, GSVA, and pan-cancer analyses. The study addressed whether an inflammation-linked expression signature could distinguish LUAD survival groups while retaining biologically interpretable associations with proliferative and immune programs.

## Materials and Methods

### Study design and public datasets

This study was a retrospective secondary analysis of de-identified, publicly available bulk transcriptomic and clinical data. The analytical workflow comprised tumor-control differential expression, inflammation-related consensus clustering, functional enrichment, survival modeling, immune-cell enrichment, pathway analysis, and pan-cancer contextualization. No new biospecimens were collected.

Expression and clinical data were obtained from the Gene Expression Omnibus (GEO) [12], The Cancer Genome Atlas (TCGA) [13], and the Genotype-Tissue Expression (GTEx) project [14]. The GEO resources were GSE11969 [15], GSE31210 [16], GSE30219 [17], and GSE40791 [18]. TCGA-LUAD and GSE31210 were designated as model-development resources; GSE11969, GSE30219, and GSE40791 were used as external expression or validation resources. After exclusion of cases without complete expression or overall-survival data, the model-development dataset contained 730 cases and was divided randomly in a 7:3 ratio into a training set (n=502) and an internal-validation set (n=228). The external survival-validation cohort consisted of 85 LUAD cases from GSE30219.

### Expression matrices and inflammation-related genes

All downstream analyses used quality-controlled, normalized gene-expression matrices assembled for the designated cohorts. Expression values entering the prognostic model were standardized before score calculation. Inflammation-associated genes were retrieved from GeneCards [19]; 596 genes were retained as the predefined inflammation-related feature set.

### Differential-expression and functional-enrichment analyses

Differential expression between LUAD and control lung samples was evaluated with limma version 3.50.0 [20]. Genes with an absolute log2 fold change greater than 1 and a Benjamini-Hochberg adjusted P value below 0.05 were classified as differentially expressed [21]. Gene Ontology (GO) categories [22] and Kyoto Encyclopedia of Genes and Genomes (KEGG) pathways [23] were evaluated with clusterProfiler version 4.2.2 [24]. Enrichment results with P<0.05 were retained.

### Inflammation-associated consensus clustering

The TCGA-LUAD expression matrix was clustered on the basis of the 596 inflammation-related genes using ConsensusClusterPlus version 1.58.0 [25]. Consensus clustering was repeated 1,000 times, and a two-cluster solution (k=2) was selected from the consensus matrix and cumulative-distribution-function profiles. Differential expression between Cluster 1 and Cluster 2 used an absolute log2 fold change greater than 1 and P<0.05.

### Identification of candidate genes and prognostic model

Inflammation-related candidate genes were defined as the intersection of LUAD-versus-control differentially expressed genes and subtype-associated differentially expressed genes. Univariate Cox proportional-hazards regression was used to test each intersecting gene for association with overall survival; genes with P<=0.05 entered penalized modeling. LASSO-Cox regression was performed with glmnet [26,27], using the minimum cross-validated partial-likelihood deviance to select the penalty. The random seed was 9725. The fitted model retained five genes: *CHRDL1, FDCSP, CXCL13, CYP4B1*, and *S100P*.

For each patient, the risk score was calculated as the sum of the standardized expression value of each retained gene multiplied by its fitted LASSO-Cox coefficient: risk score = Σ (β_i_ × z_i_), where z_i denotes standardized gene expression and beta_i denotes the fitted coefficient. Each analyzed cohort was separated into high- and low-risk groups at its median risk score. Overall survival was compared with Kaplan-Meier curves and the two-sided log-rank test. Time-dependent receiver operating characteristic curves and areas under the curve (AUCs) were calculated at 1, 3, and 5 years [28].

### Immune-cell and pathway analyses

Twenty-eight immune-cell gene signatures were obtained through TISIDB [33] and the underlying pan-cancer immunogenomic framework [34]. Single-sample gene set enrichment analysis was used to quantify relative immune-cell enrichment in each LUAD tumor. High- and low-risk groups were compared with the Wilcoxon rank-sum test, and immune-score relationships were summarized by a correlation matrix. Visualizations were generated with ggplot2 version 3.3.6 [35].

GSEA was performed with clusterProfiler on all genes ranked by the log2 fold change for high-risk versus low-risk tumors [24,29]. The c2.cp.kegg.v7.5.1.symbols collection from the Molecular Signatures Database (MSigDB) was used as the reference [30]. Each analysis used 1,000 permutations, and gene sets with P<0.05 were retained. Positive normalized enrichment scores indicated enrichment toward the high-risk end of the ranked list, whereas negative scores indicated enrichment toward the low-risk end.

GSVA version 1.42.0 was applied to MSigDB Hallmark gene sets [31,32]. Pathway scores were compared between risk groups with the Wilcoxon rank-sum test and visualized with pheatmap version 1.0.12.

### Pan-cancer analysis

TCGA expression and clinical data for 33 cancer types were obtained with TCGAbiolinks version 2.25.0 [36]. Tumor-versus-normal expression differences for signature genes were assessed with the Wilcoxon rank-sum test. Univariate Cox regression was used to evaluate gene-expression associations with overall survival. Immune-cell enrichment scores were calculated by single-sample GSEA, and gene-expression/immune-score correlations were summarized across cancer types.

### Statistical analysis and ethics

Continuous variables were compared with the Wilcoxon rank-sum test. Categorical proportions were compared with the chi-square test or Fisher exact test, as appropriate. Kaplan-Meier curves were fitted with survminer, and survival differences were assessed with the log-rank test. Unless otherwise specified, tests were two-sided and P<0.05 was considered statistically significant. Analyses were performed in R version 4.1.2. Because only de-identified public datasets were analyzed and no new participant recruitment or specimen collection occurred, the study did not involve direct human-subject intervention.

## Results

### Public transcriptomic resources and analysis framework

The study assembled six public resources spanning LUAD tumor tissue, adjacent or non-neoplastic lung tissue, and GTEx normal lung (Table 1). The model-development dataset comprised 730 cases, including 502 training cases and 228 internal-validation cases. External survival validation used 85 GSE30219 cases. The analysis integrated tumor-control expression differences, inflammation-defined tumor heterogeneity, survival modeling, immune-state analysis, pathway-level interpretation, and pan-cancer comparison.

**Table 1.** Public transcriptomic resources and analytical roles.

| Dataset | Repository/platform | Cases used | Control samples | Primary role |
| --- | --- | --- | --- | --- |
| GSE11969 | GEO / GPL7015 | 158 tumor samples; 149 with survival data | 5 normal | External expression and survival resource |
| GSE30219 | GEO / GPL570 | 85 LUAD cases used for external survival validation | Tumor-normal expression panel included | External validation |
| GSE31210 | GEO / GPL570 | 226 LUAD samples; 226 with survival data | 20 normal | Model-development resource |
| GSE40791 | GEO / GPL570 | 94 LUAD samples | 100 non-neoplastic lung | External expression resource |
| TCGA-LUAD | TCGA / RNA sequencing | 513 tumor samples; 513 with survival data | 59 adjacent normal | Discovery and model-development resource |
| GTEx lung | GTEx / RNA sequencing | Not applicable | 288 normal lung | Normal-lung reference |
| Prognostic dataset | Integrated development resource | 730 cases with complete prognosis; 502 training and 228 internal validation | Not applicable | Risk-score construction and internal validation |

Counts represent the analytic sample numbers used in the uploaded analysis. OS, overall survival; TCGA, The Cancer Genome Atlas; GTEx, Genotype-Tissue Expression; GEO, Gene Expression Omnibus.

**Table 2.**
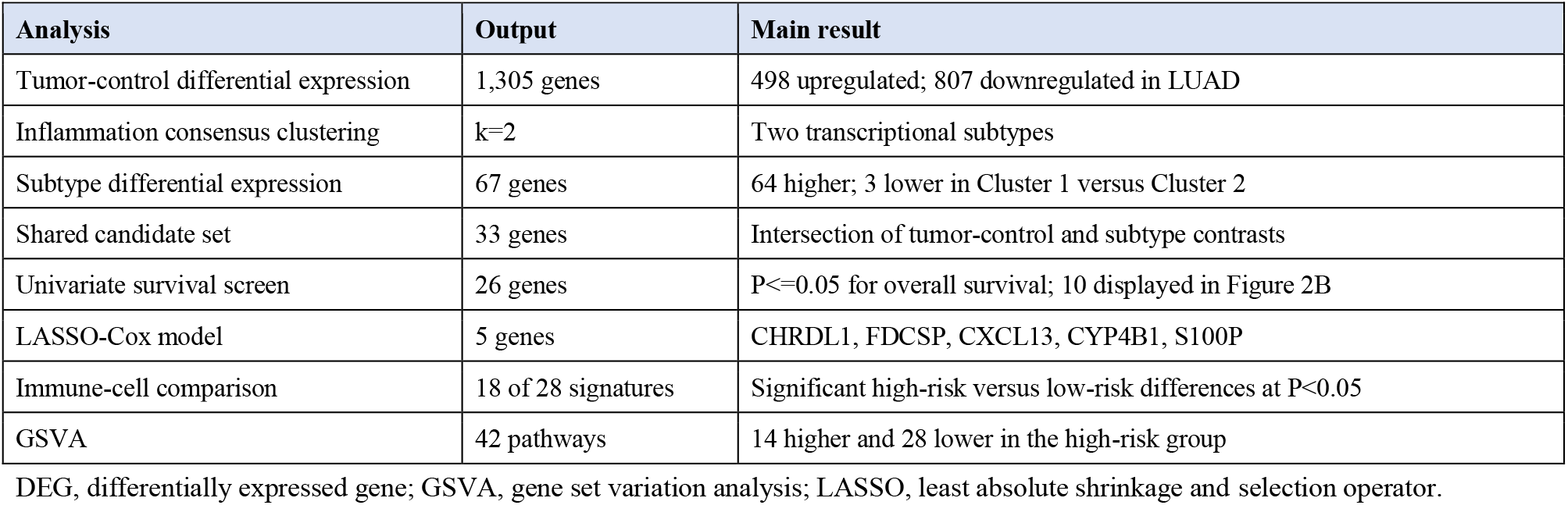
Summary of principal analytical results.

**Table 3.** Time-dependent discrimination of the five-gene risk score.

| Cohort | n | 1-year AUC | 3-year AUC | 5-year AUC | Survival comparison |
| --- | --- | --- | --- | --- | --- |
| Training | 502 | 0.6625 | 0.6581 | 0.6658 | High-risk survival was poorer; log-rank P<0.05 |
| Internal validation | 228 | 0.7422 | 0.6537 | 0.6761 | High-risk survival was poorer; log-rank P<0.05 |
| External validation (GSE30219) | 85 | 0.6560 | 0.6387 | 0.6753 | High-risk survival was poorer; log-rank P<0.05 |

AUC, area under the time-dependent receiver operating characteristic curve.

**Table 4.**
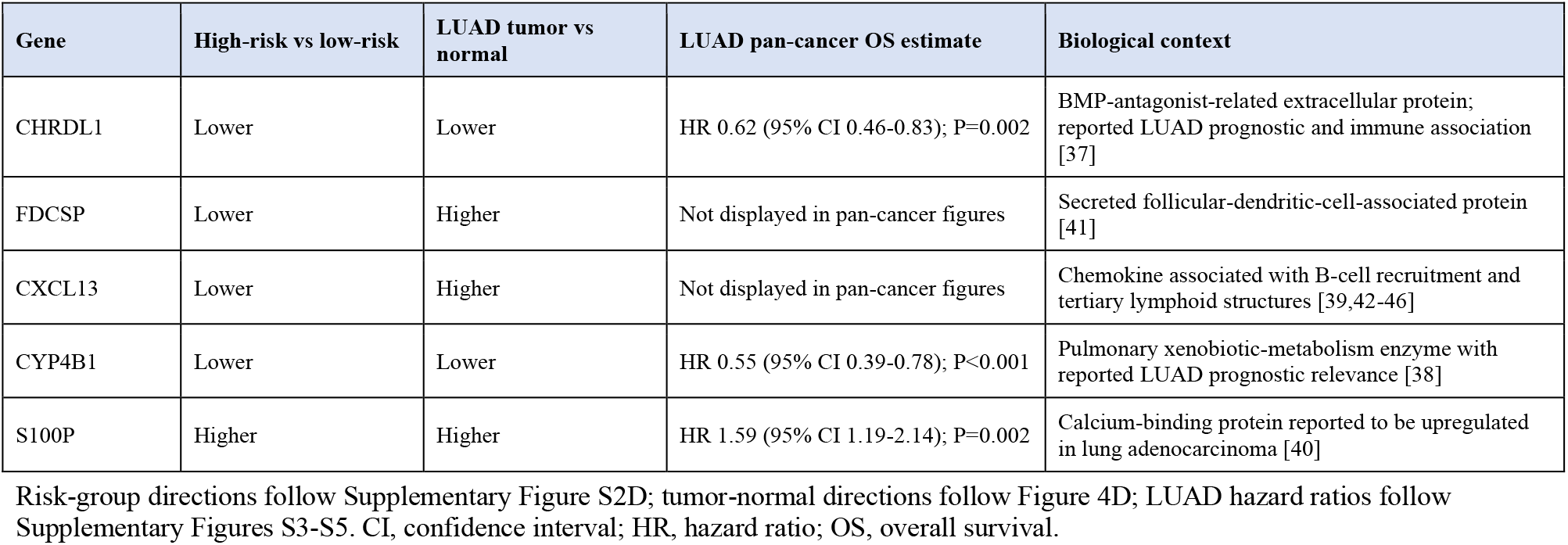
Expression and survival context of the five-gene signature.

### LUAD and control tissues showed broad expression differences

The LUAD-versus-control analysis identified 1,305 differentially expressed genes, including 498 genes upregulated and 807 genes downregulated in LUAD (Supplementary Figure S1A). *OCIAD2, GOLM1, PYCR1, ALDH18A1*, and *B3GNT3* were among the most strongly upregulated genes, whereas *FENDRR, STARD8, LIMS2, TAL1*, and *ACVRL1* were among the most strongly downregulated genes (Supplementary Figure S1B,C).

GO enrichment emphasized cell-substrate adhesion, external encapsulating-structure organization, extracellular-matrix organization, collagen-containing extracellular matrix, cell-cell junctions, extracellular-matrix structural constituents, glycosaminoglycan binding, and heparin binding (Supplementary Figure S1D). KEGG enrichment included focal adhesion, cytoskeleton in muscle cells, cell cycle, ECM-receptor interaction, PI3K-Akt signaling, DNA replication, non-homologous end joining, Fanconi anemia, leukocyte transendothelial migration, and selected metabolic pathways (Supplementary Figure S1E). These results connected LUAD-associated expression changes to extracellular-structure remodeling, adhesion, proliferation, and genome-maintenance programs.

### Inflammation-related expression separated LUAD into two subtypes

Consensus clustering of TCGA-LUAD tumors on the basis of inflammation-related gene expression supported a two-cluster solution (Figure 1A,B). Differential-expression analysis between Cluster 1 and Cluster 2 identified 67 subtype-associated genes. Sixty-four genes were higher and three genes were lower in Cluster 1 relative to Cluster 2 (Figure 1C). *MNDA, PTPRC, MS4A6A, HLA-DPB1*, and *CD52* were prominent higher-expression genes, whereas *FGL1, FGB*, and *S100P* were the three lower-expression genes (Figure 1D,E).

**Figure 1.**
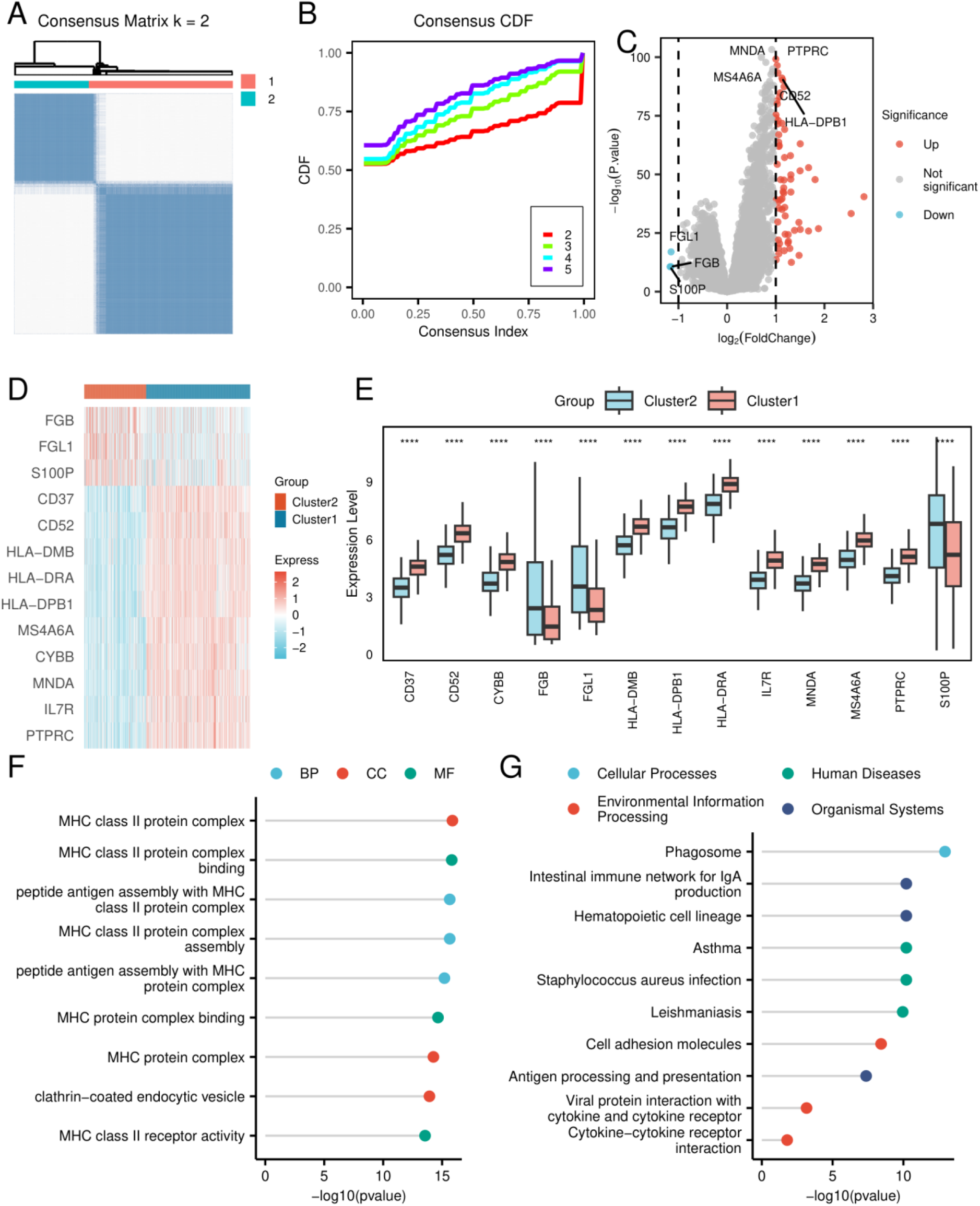
Inflammation-associated consensus clustering and subtype differences in LUAD. (A) Consensus matrix for k=2. (B) Consensus cumulative-distribution-function curves across candidate cluster numbers. (C) Volcano plot comparing Cluster 1 with Cluster 2. (D) Heatmap of selected subtype-associated genes. (E) Boxplots of displayed genes between clusters. (F) GO enrichment of the 67 subtype-associated genes. (G) KEGG enrichment of subtype-associated genes. The comparison identified 64 genes with higher expression and 3 genes with lower expression in Cluster 1 relative to Cluster 2.

Subtype-associated genes were enriched for MHC class II protein-complex assembly, peptide-antigen assembly with MHC class II, MHC class II complex binding and receptor activity, and MHC protein complexes (Figure 1F). KEGG terms included phagosome, efferocytosis, ferroptosis, cell adhesion molecules, cytokine-cytokine receptor interaction, antigen processing and presentation, hematopoietic cell lineage, and the intestinal immune network for IgA production (Figure 1G). Thus, the two subtypes differed predominantly in antigen-presentation and immune-related transcriptional programs.

### Thirty-three genes linked tumor dysregulation to inflammation-defined heterogeneity

Intersecting the 1,305 tumor-control differentially expressed genes with the 67 subtype-associated genes yielded 33 inflammation-related candidate genes (Figure 2A). GO terms included regulation of cell-cell adhesion, humoral immune response, regulation of T-cell activation, lamellar bodies, multivesicular bodies, collagen-containing extracellular matrix, water-channel activity, water transmembrane-transporter activity, and cargo-receptor activity (Figure 2C). The KEGG panel highlighted phagosome and complement/coagulation cascades (Figure 2D).

**Figure 2.**
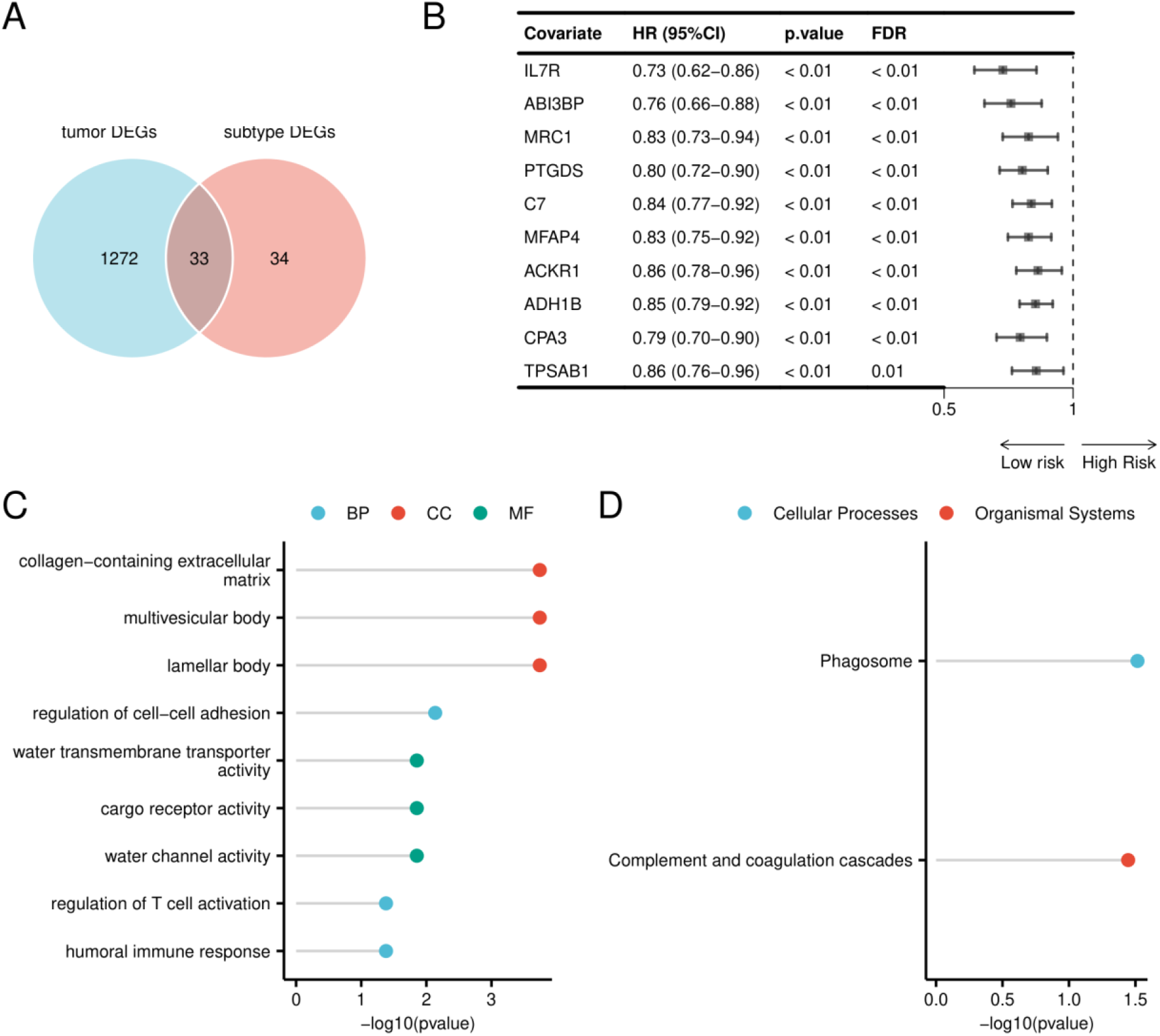
Identification and functional characterization of inflammation-related candidate genes. (A) Venn diagram showing 1,272 tumor-control-specific genes, 34 subtype-specific genes, and 33 shared genes. (B) Forest plot of ten displayed overall-survival-associated genes from univariate Cox regression; points indicate HRs and horizontal lines indicate 95% CIs. (C) GO enrichment of the 33 shared genes. (D) KEGG enrichment of the shared genes.

Univariate Cox analysis identified 26 of the 33 genes as overall-survival associated. Figure 2B displayed ten protective associations: *IL7R* (hazard ratio [HR] 0.73, 95% confidence interval [CI] 0.62-0.86), *ABI3BP* (HR 0.76, 95% CI 0.66-0.88), *MRC1* (HR 0.83, 95% CI 0.73-0.94), *PTGDS* (HR 0.80, 95% CI 0.72-0.90), *C7* (HR 0.84, 95% CI 0.77-0.92), *MFAP4* (HR 0.83, 95% CI 0.75-0.92), *ACKR1* (HR 0.86, 95% CI 0.78-0.96), *ADH1B* (HR 0.85, 95% CI 0.79-0.92), *CPA3* (HR 0.79, 95% CI 0.70-0.90), and *TPSAB1* (HR 0.86, 95% CI 0.76-0.96). The displayed associations had P<0.01 and false-discovery-rate values at or below 0.01.

### A five-gene score stratified overall survival in the training set

LASSO-Cox regression in the 502-case training set retained *CHRDL1, FDCSP, CXCL13, CYP4B1*, and *S100P* (Figure 3A,B). The median risk score separated patients into high- and low-risk groups. Mortality increased across the ordered risk-score distribution, and the five-gene heatmap showed coordinated expression changes across the risk continuum (Figure 3C).

**Figure 3.**
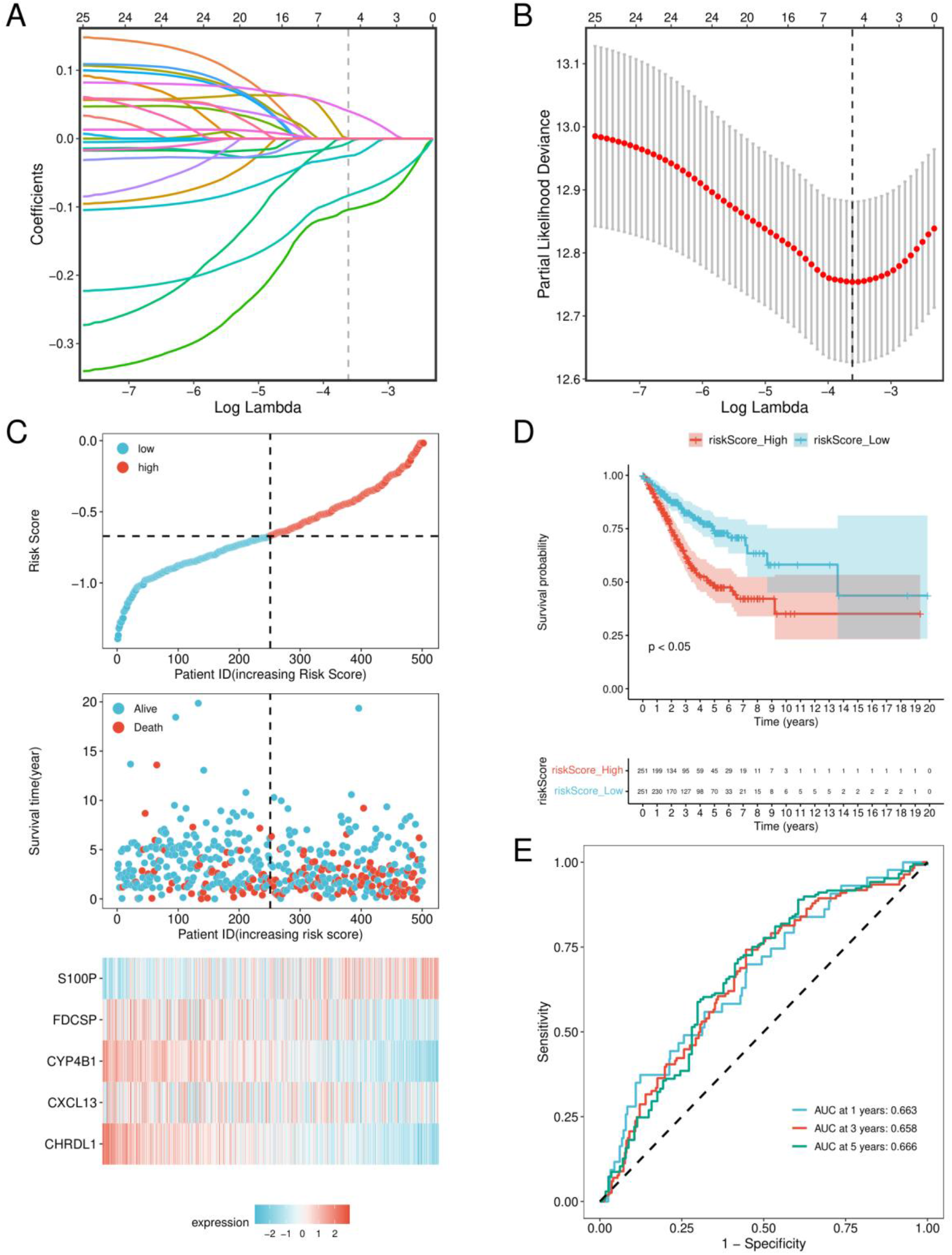
Derivation of the five-gene LUAD risk score in the training cohort. (A) LASSO-Cox coefficient trajectories as a function of log(lambda). (B) Cross-validated partial-likelihood deviance used to select the penalty at the minimum criterion. (C) Risk-score distribution, survival-status distribution, and expression heatmap for *CHRDL1, FDCSP, CXCL13, CYP4B1*, and *S100P* in the training set (n=502). (D) Kaplan-Meier overall-survival curves for high- and low-risk groups, compared with the two-sided log-rank test. (E) Time-dependent receiver operating characteristic curves at 1, 3, and 5 years; AUCs were 0.6625, 0.6581, and 0.6658, respectively.

The high-risk group had poorer overall survival than the low-risk group by log-rank testing (P<0.05; Figure 3D). Time-dependent AUCs were 0.6625 at 1 year, 0.6581 at 3 years, and 0.6658 at 5 years (Figure 3E), indicating moderate discrimination in the training set.

### Internal validation reproduced survival separation

In the 228-case internal-validation set, patients were divided at the median risk score. The ordered risk plot again showed increased mortality at higher scores (Supplementary Figure S2A). The high-risk group had poorer overall survival than the low-risk group (log-rank P<0.05; Supplementary Figure S2B). The 1-, 3-, and 5-year AUCs were 0.7422, 0.6537, and 0.6761, respectively (Supplementary Figure S2C). *CHRDL1, CXCL13, CYP4B1*, and *FDCSP* were lower in the high-risk group, whereas *S100P* was higher (Supplementary Figure S2D).

### External validation in GSE30219 confirmed risk-group differences

External validation used 85 LUAD cases from GSE30219. The risk distribution and survival-status panel showed increased deaths toward the high-risk end of the score (Figure 4A). High-risk patients had poorer overall survival than low-risk patients (log-rank P<0.05; Figure 4B). The 1-, 3-, and 5-year AUCs were 0.6560, 0.6387, and 0.6753, respectively (Figure 4C). In the accompanying tumor-control expression panel, *CHRDL1* and *CYP4B1* were lower in LUAD tumors, whereas *CXCL13, FDCSP*, and *S100P* were higher (Figure 4D).

**Figure 4.**
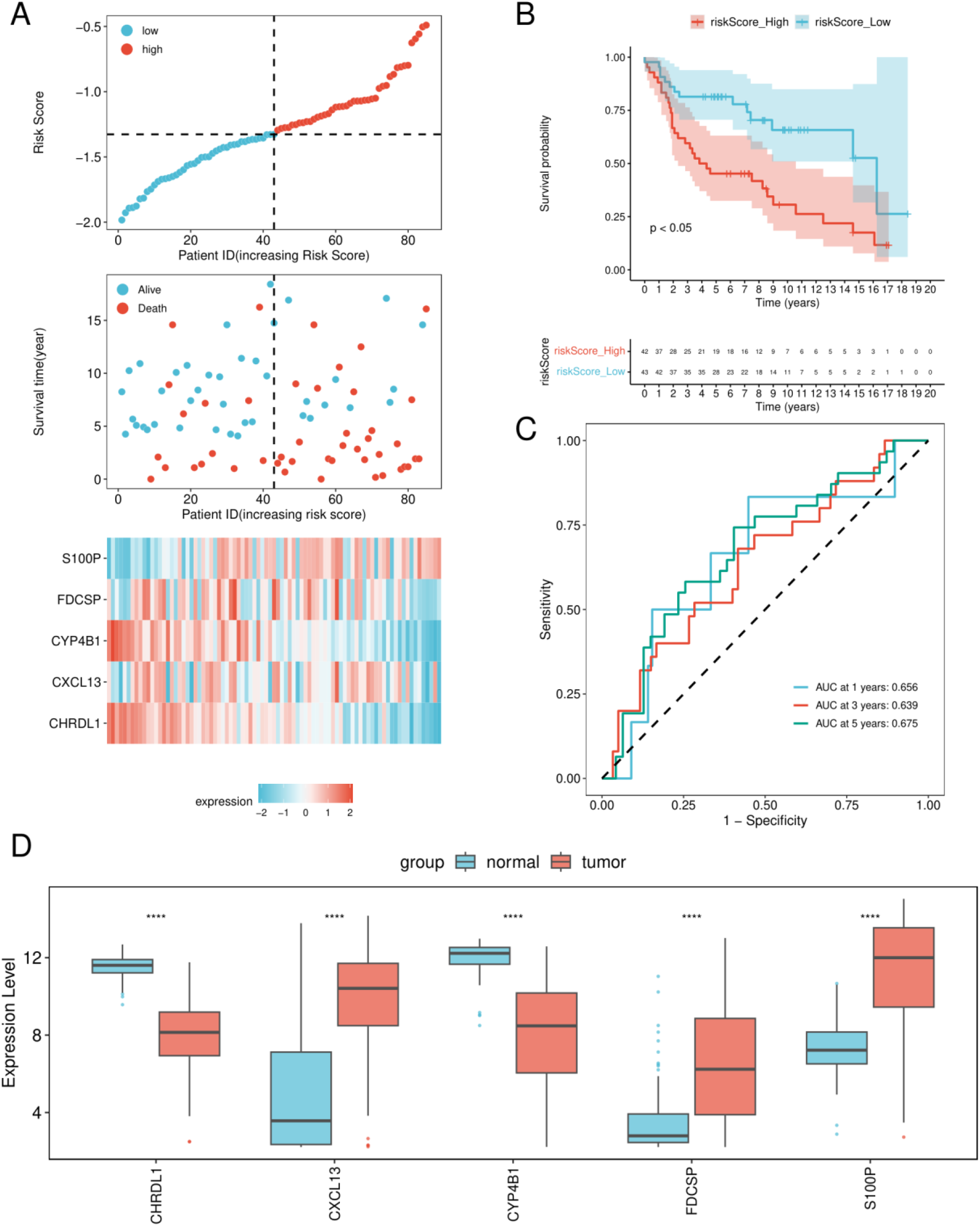
External validation of the five-gene risk score in GSE30219. (A) Risk-score distribution, survival-status distribution, and five-gene expression heatmap for the 85-case external-validation cohort. (B) Kaplan-Meier overall-survival curves for high- and low-risk groups. (C) Time-dependent receiver operating characteristic curves at 1, 3, and 5 years; AUCs were 0.6560, 0.6387, and 0.6753, respectively. (D) Expression of *CHRDL1, CXCL13, CYP4B1, FDCSP*, and *S100P* in normal and LUAD tumor samples. Significance symbols: ****P<0.0001, ***P<0.001, **P<0.01, and *P<0.05.

### Risk groups differed in immune-cell enrichment

Relative enrichment scores were estimated for 28 immune-cell signatures across LUAD tumors (Figure 5A). Significant high-risk versus low-risk differences were observed for effector-memory CD8 T cells, activated CD4 T cells, effector-memory CD4 T cells, follicular-helper T cells, gamma-delta T cells, type 2 helper T cells, activated B cells, immature B cells, memory B cells, CD56bright natural-killer cells, CD56dim natural-killer cells, myeloid-derived suppressor cells, natural-killer T cells, plasmacytoid dendritic cells, macrophages, eosinophils, mast cells, and neutrophils (P<0.05; Figure 5C). Correlation analysis showed broad positive relationships among many immune signatures (Figure 5B), indicating coordinated variation in the immune composition represented by bulk-tissue expression.

**Figure 5.**
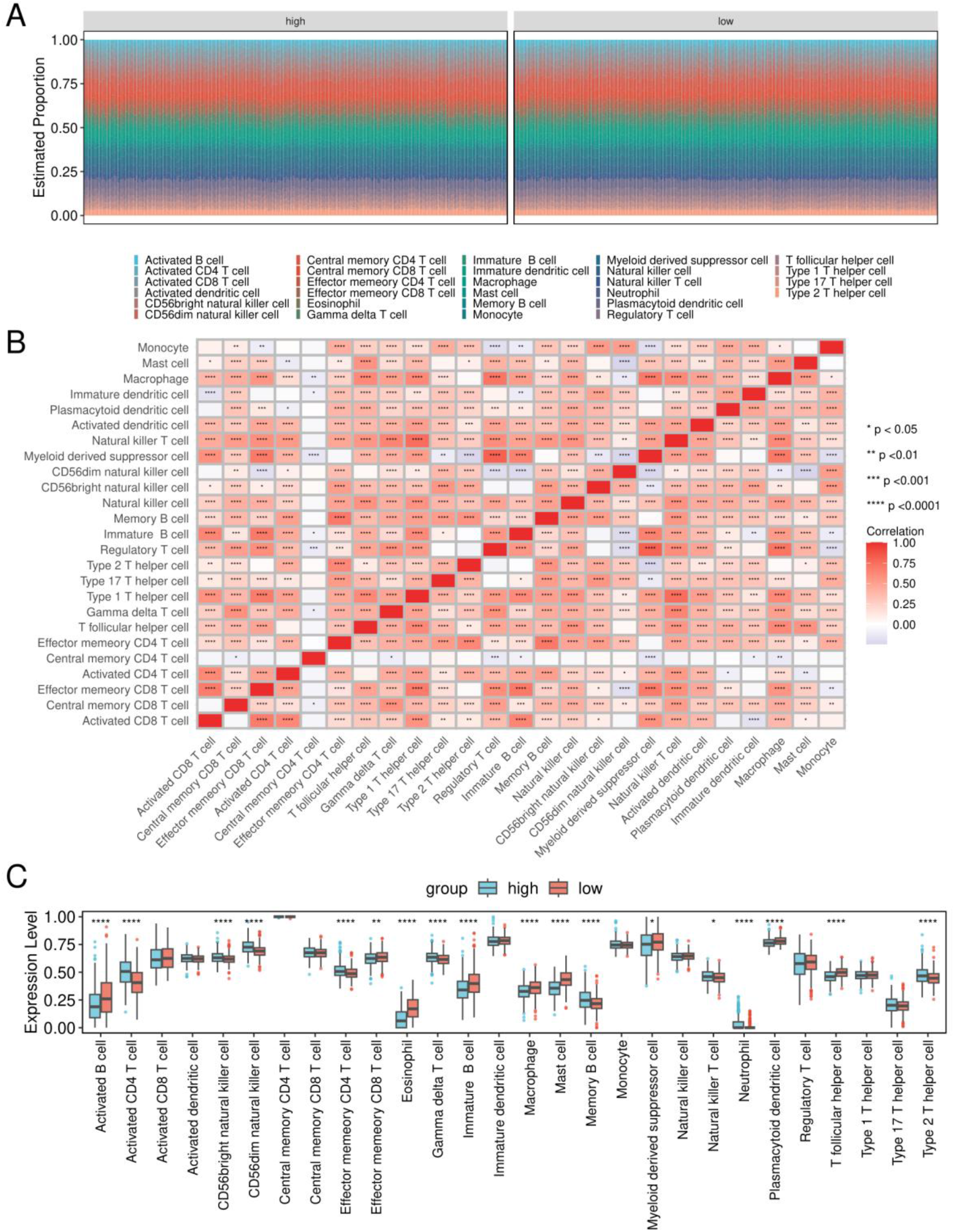
Immune-cell enrichment in LUAD risk groups. (A) Relative enrichment of 28 immune-cell signatures across LUAD samples. (B) Correlation matrix of immune-cell enrichment scores. (C) Comparison of immune-cell enrichment scores between high- and low-risk groups. Values represent transcriptome-derived enrichment scores rather than direct cell counts. Significance symbols: ****P<0.0001, ***P<0.001, **P<0.01, and *P<0.05.

### Proliferation and DNA-maintenance programs distinguished risk groups

GSEA identified positive enrichment toward the high-risk end of the ranked expression profile for CELL CYCLE (NES=2.67; adjusted P=1.42×10^−8^), DNA REPLICATION (NES=2.52; adjusted P=2.52×10^−7^), and MISMATCH REPAIR (NES=2.20; adjusted P=1.77×10^−4^) (Figure 6A-C). Negative enrichment toward the low-risk end was observed for CELL ADHESION MOLECULES CAMS (NES=-2.25; adjusted P=1.42×10^−8^), VIRAL MYOCARDITIS (NES=-2.24; adjusted P=2.81×10^−7^), and ASTHMA (NES=-2.23; adjusted P=8.03×10^−6^) (Figure 6D-F).

**Figure 6.**
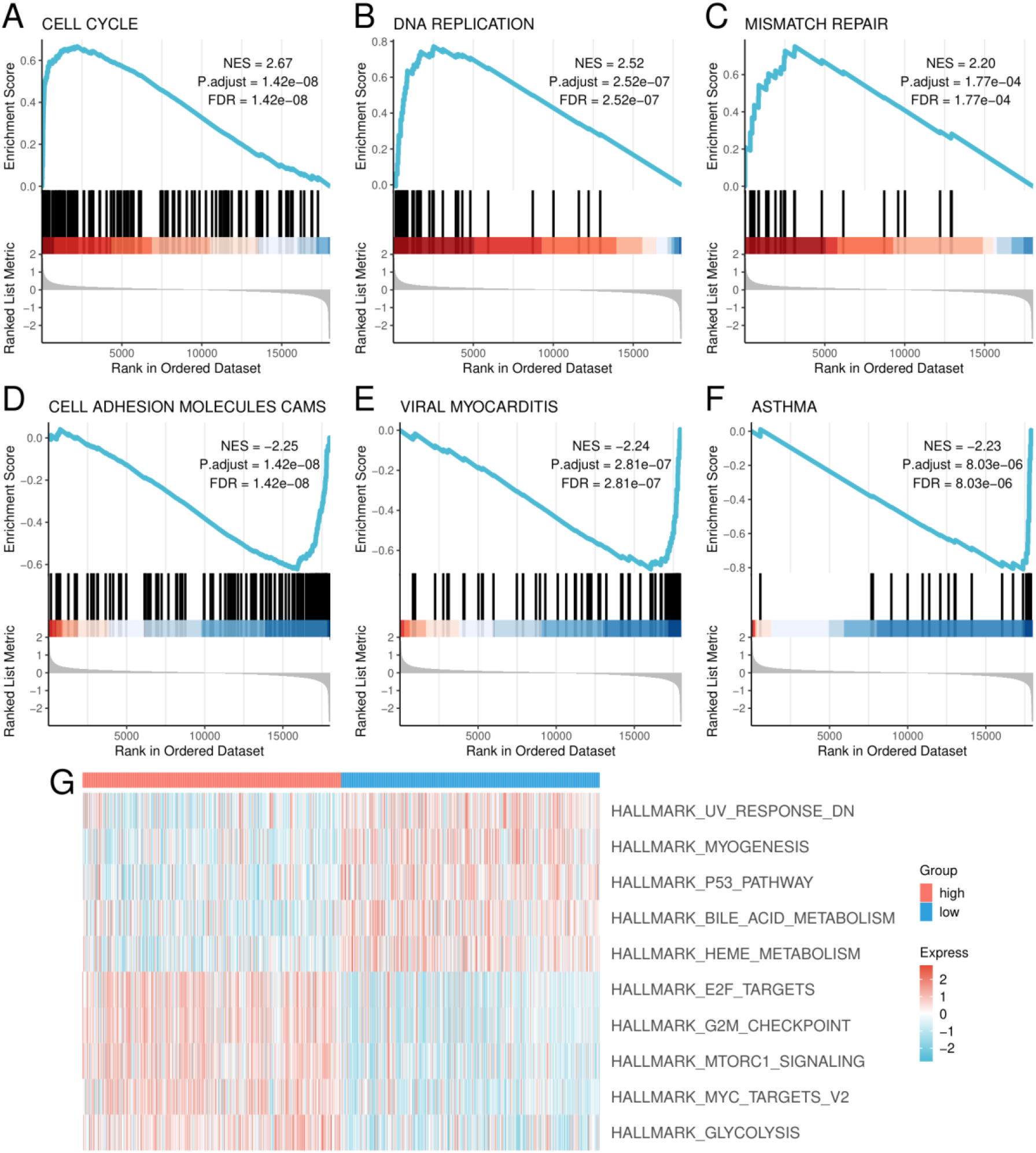
GSEA and GSVA of high- and low-risk LUAD groups. (A-F) GSEA plots for cell cycle, DNA replication, mismatch repair, cell adhesion molecules, viral myocarditis, and asthma. Positive NES values indicate enrichment toward the high-risk end of the ranked list; negative values indicate enrichment toward the low-risk end. (G) Heatmap of selected Hallmark GSVA scores. NES, normalized enrichment score; GSEA, gene set enrichment analysis; GSVA, gene set variation analysis.

Hallmark GSVA identified 14 pathways with higher scores in the high-risk group, including G2M_CHECKPOINT, E2F_TARGETS, MYC_TARGETS_V2, MTORC1_SIGNALING, and GLYCOLYSIS.

Twenty-eight pathways had lower scores in the high-risk group, including HEME_METABOLISM, MYOGENESIS, P53_PATHWAY, UV_RESPONSE_DN, and BILE_ACID_METABOLISM (Figure 6G).

Together, these analyses associated higher risk with stronger proliferative, DNA-replication, and metabolic programs and with altered adhesion and immune-related states.

### Pan-cancer analyses contextualized CHRDL1, CYP4B1, and S100P

Pan-cancer analysis showed heterogeneous tumor-normal expression, survival associations, and immune-score correlations for *CHRDL1, CYP4B1*, and *S100P* across TCGA cancer types. In LUAD, higher *CHRDL1* expression was associated with lower mortality risk (HR 0.62, 95% CI 0.46-0.83; P=0.002), and *CHRDL1* expression was lower in tumor than in normal lung (Supplementary Figure S3).

Higher *CYP4B1* expression was also associated with lower mortality risk in LUAD (HR 0.55, 95% CI 0.39-0.78; P<0.001), and *CYP4B1* was lower in LUAD tumor tissue than in normal lung (Supplementary Figure S4).

*S100P* showed the opposite LUAD survival direction: higher expression was associated with greater mortality risk (HR 1.59, 95% CI 1.19-2.14; P=0.002), and *S100P* expression was higher in LUAD tumor tissue than in normal lung (Supplementary Figure S5). The cross-cancer heatmaps showed that immune associations varied by cancer type, indicating context-dependent relationships rather than a uniform pan-cancer effect.

## Discussion

This integrative analysis identified two inflammation-associated LUAD expression states and derived a five-gene score comprising *CHRDL1, FDCSP, CXCL13, CYP4B1*, and *S100P*. The score separated high- and low-risk groups in training, internal-validation, and external-validation cohorts. Discrimination was moderate, with AUCs between approximately 0.64 and 0.74, but the biological analyses were coherent: higher risk was associated with stronger cell-cycle, DNA-replication, G2M, E2F, MYC, mTORC1, and glycolytic programs, together with broad differences in immune-cell enrichment.

The tumor-control analysis emphasized extracellular-matrix organization, focal adhesion, cell-cell junctions, PI3K-Akt signaling, and cell-cycle programs. These processes are consistent with the reciprocal interactions among cancer cells, inflammatory mediators, stromal cells, and extracellular matrix that shape tumor progression [4-6,10,11]. The inflammation-defined subtype contrast added an immunologic layer, with strong representation of MHC class II assembly, antigen processing and presentation, phagosome, cytokine interactions, and hematopoietic-lineage programs. Intersecting the two contrasts therefore prioritized genes that were both altered in LUAD tissue and variable across inflammation-associated tumor states.

The five selected genes represented distinct biological components of the signature. Higher *CHRDL1* and *CYP4B1* expression aligned with the lower-risk state, and independent LUAD studies have linked reduced expression of these genes to adverse clinical or immune features [37,38]. *S100P* showed the opposite pattern: it was higher in high-risk tumors, higher in LUAD than normal lung, and associated with increased mortality risk in the LUAD pan-cancer analysis. Experimental work has documented *S100P* upregulation and regulation in human lung adenocarcinoma [40]. *FDCSP* was originally characterized as a secreted follicular-dendritic-cell-associated protein [41], making its inclusion compatible with the inflammation-focused feature-selection strategy, although its direct function in LUAD remains uncertain.

*CXCL13* provides a plausible link between the lower-risk state and organized humoral immunity. In LUAD treated with immune-checkpoint inhibitors, higher *CXCL13* expression has been associated with immune-response genes, tertiary lymphoid structure signatures, and favorable outcomes [39]. In lung cancer, intratumoral tertiary lymphoid structures and their B-cell compartments have been associated with prolonged survival and protective immune states [42,43]. Related studies across melanoma and sarcoma further support the relationship among B cells, tertiary lymphoid structures, survival, and checkpoint-blockade response [44-46]. The present analysis did not test immunotherapy response, so *CXCL13* should be interpreted as an immune-context marker rather than a treatment-predictive biomarker.

The risk groups differed across multiple T-cell, B-cell, natural-killer-cell, myeloid, dendritic-cell, macrophage, and granulocyte signatures. The broad covariance among these scores is consistent with coordinated immune ecosystems rather than isolated cell lineages [47]. However, expression-derived immune scores depend on the signatures and computational framework used. Established approaches such as CIBERSORT, xCell, MCP-counter, and quanTIseq illustrate both the utility and the method dependence of bulk-transcriptome immune inference [48-51]. The present values should therefore be interpreted as relative enrichment states, not direct histologic cell counts or proof of immune-cell function.

The GSEA and GSVA results indicate that the risk score captured more than a single immune axis. Positive enrichment of cell cycle, DNA replication, mismatch repair, G2M checkpoint, E2F targets, MYC targets, mTORC1 signaling, and glycolysis suggests a proliferative and metabolically active high-risk state. Negative enrichment of cell-adhesion and selected immune-related pathways, together with lower scores for heme metabolism, myogenesis, p53 pathway, ultraviolet-response, and bile-acid-metabolism programs, suggests broader tissue-state remodeling. These pathway-level patterns provide biological context but do not establish that any individual pathway causally determines survival.

The model showed reproducible direction of survival separation, but its AUCs were insufficient for stand-alone clinical decision-making. A clinically deployable prognostic tool would require a fully locked preprocessing pipeline, fixed model coefficients and threshold, calibration, confidence intervals, comparison with established clinicopathologic variables, net-benefit analysis, and validation in independent prospective or temporally separated cohorts. These requirements are consistent with TRIPOD reporting principles and PROBAST risk-of-bias domains [52,53]. The current score is best viewed as an exploratory transcriptomic stratifier and a basis for subsequent validation.

### Limitations

- The study was retrospective and relied on public cohorts generated on different transcriptomic platforms. Platform heterogeneity and cohort-specific preprocessing may influence differential expression, risk scores, and immune enrichment.
- The external survival-validation cohort contained 85 cases, limiting precision and subgroup analysis. Additional validation across institutions, ethnic groups, disease stages, and treatment contexts is needed.
- The risk score was evaluated primarily by Kaplan-Meier separation and time-dependent AUC. Multivariable adjustment for age, sex, smoking status, stage, molecular alterations, treatment, and tumor purity was not included in the available analysis.
- Median-based risk stratification is suitable for exploratory comparison but does not define a clinically transportable threshold. Calibration, Brier scores, decision curves, and repeated resampling were not part of the reported analysis.
- Bulk-transcriptome immune scores cannot distinguish whether a gene signal arises from malignant cells, stromal cells, or infiltrating immune cells. Single-cell, spatial, pathology-based, and protein-level validation would be required to establish cellular origin.
- No experimental perturbation or prospective clinical validation was performed. The observed gene, pathway, immune, and survival relationships are associations and do not establish causality.

### Future validation

The next computational step is to test the five-gene score in independent LUAD cohorts using a fixed transformation, coefficient vector, and threshold, followed by multivariable Cox modeling, proportional-hazards diagnostics, calibration, and bootstrap or repeated-cross-validation assessment. Biological validation should quantify the five transcripts in an independent LUAD series, evaluate protein localization where suitable, and use single-cell or spatial transcriptomics to identify the cellular sources of each signal. Functional experiments should prioritize genes whose expression, survival direction, and cell-of-origin relationships are reproduced across these orthogonal analyses.

## Conclusions

An integrated analysis of public LUAD transcriptomes identified two inflammation-associated expression subtypes and a five-gene score comprising *CHRDL1, FDCSP, CXCL13, CYP4B1*, and *S100P*. The score separated overall-survival groups in training, internal-validation, and external-validation cohorts and was associated with coordinated proliferative, metabolic, adhesion, and immune transcriptional programs. These findings establish a testable inflammation-linked stratification framework. Independent clinical adjustment, locked-model validation, and experimental confirmation are required before the score can be considered for clinical use.

## Supplementary Materials

Supplementary Figures S1-S5 are provided in a separate file. Supplementary Figure S1 presents the LUAD-versus-control differential-expression and enrichment analysis; Supplementary Figure S2 presents internal validation of the five-gene score; and Supplementary Figures S3-S5 present pan-cancer analyses of CHRDL1, CYP4B1, and S100P, respectively.

## Notes

### Competing Interest Statement

The authors have declared no competing interest.

## References

1. Sung H, Filho AM, Laversanne M, Ferlay J, Siegel RL, Soerjomataram I, et al. Global cancer statistics 2024: GLOBOCAN estimates of incidence and mortality worldwide for 34 cancers in 186 countries. CA Cancer J Clin. 2026;76(4):e70090. doi: 10.3322/caac.70090. PMID:42417444.

2. Herbst RS, Morgensztern D, Boshoff C. The biology and management of non-small cell lung cancer. Nature. 2018;553(7689):446–454. doi: 10.1038/nature25183. PMID:29364287.

3. Cancer Genome Atlas Research Network. Comprehensive molecular profiling of lung adenocarcinoma. Nature. 2014;511(7511):543–550. doi: 10.1038/nature13385. PMID:25079552.

4. Coussens LM, Werb Z. Inflammation and cancer. Nature. 2002;420(6917):860–867. doi: 10.1038/nature01322. PMID:12490959.

5. Mantovani A, Allavena P, Sica A, Balkwill F. Cancer-related inflammation. Nature. 2008;454(7203):436–444. doi: 10.1038/nature07205. PMID:18650914.

6. Grivennikov SI, Greten FR, Karin M. Immunity, inflammation, and cancer. Cell. 2010;140(6):883–899. doi: 10.1016/j.cell.2010.01.025. PMID:20303878.

7. Binnewies M, Roberts EW, Kersten K, Chan V, Fearon DF, Merad M, et al. Understanding the tumor immune microenvironment (TIME) for effective therapy. Nat Med. 2018;24(5):541–550. doi: 10.1038/s41591-018-0014-x. PMID:29686425.

8. Thorsson V, Gibbs DL, Brown SD, Wolf D, Bortone DS, Ou Yang TH, et al. The Immune Landscape of Cancer. Immunity. 2018;48(4):812–830.e14. doi: 10.1016/j.immuni.2018.03.023. PMID:29628290.

9. Fridman WH, Pages F, Sautes-Fridman C, Galon J. The immune contexture in human tumours: impact on clinical outcome. Nat Rev Cancer. 2012;12(4):298–306. doi: 10.1038/nrc3245. PMID:22419253.

10. Joyce JA, Pollard JW. Microenvironmental regulation of metastasis. Nat Rev Cancer. 2009;9(4):239–252. doi: 10.1038/nrc2618. PMID:19279573.

11. Pickup MW, Mouw JK, Weaver VM. The extracellular matrix modulates the hallmarks of cancer. EMBO Rep. 2014;15(12):1243–1253. doi: 10.15252/embr.201439246. PMID:25381661.

12. Clough E, Barrett T, Wilhite SE, Ledoux P, Evangelista C, Kim IF, et al. NCBI GEO: archive for gene expression and epigenomics data sets: 23-year update. Nucleic Acids Res. 2024;52(D1):D138–D144. doi: 10.1093/nar/gkad965. PMID:37933855.

13. Weinstein JN, Collisson EA, Mills GB, Shaw KRM, Ozenberger BA, Ellrott K, et al. The Cancer Genome Atlas Pan-Cancer analysis project. Nat Genet. 2013;45(10):1113–1120. doi: 10.1038/ng.2764. PMID:24071849.

14. GTEx Consortium. The GTEx Consortium atlas of genetic regulatory effects across human tissues. Science. 2020;369(6509):1318–1330. doi: 10.1126/science.aaz1776. PMID:32913098.

15. Takeuchi T, Tomida S, Yatabe Y, Kosaka T, Osada H, Yanagisawa K, et al. Expression profile-defined classification of lung adenocarcinoma shows close relationship with underlying major genetic changes and clinicopathologic behaviors. J Clin Oncol. 2006;24(11):1679–1688. doi: 10.1200/JCO.2005.03.8224. PMID:16549822.

16. Okayama H, Kohno T, Ishii Y, Shimada Y, Shiraishi K, Iwakawa R, et al. Identification of genes upregulated in ALK-positive and EGFR/KRAS/ALK-negative lung adenocarcinomas. Cancer Res. 2012;72(1):100–111. doi: 10.1158/0008-5472.CAN-11-1403. PMID:22080568.

17. Rousseaux S, Debernardi A, Jacquiau B, Vitte AL, Vesin A, Nagy-Mignotte H, et al. Ectopic activation of germline and placental genes identifies aggressive metastasis-prone lung cancers. Sci Transl Med. 2013;5(186):186ra66. doi: 10.1126/scitranslmed.3005723. PMID:23698379.

18. Zhang Y, Foreman O, Wigle DA, Kosari F, Vasmatzis G, Salisbury JL, et al. USP44 regulates centrosome positioning to prevent aneuploidy and suppress tumorigenesis. J Clin Invest. 2012;122(12):4362–4374. doi: 10.1172/JCI63084. PMID:23187126.

19. Stelzer G, Rosen N, Plaschkes I, Zimmerman S, Twik M, Fishilevich S, et al. The GeneCards Suite: from gene data mining to disease genome sequence analyses. Curr Protoc Bioinformatics. 2016;54:1.30.1-1.30.33. doi: 10.1002/cpbi.5. PMID:27322403.

20. Ritchie ME, Phipson B, Wu D, Hu Y, Law CW, Shi W, et al. limma powers differential expression analyses for RNA-sequencing and microarray studies. Nucleic Acids Res. 2015;43(7):e47. doi: 10.1093/nar/gkv007. PMID:25605792.

21. Benjamini Y, Hochberg Y. Controlling the false discovery rate: a practical and powerful approach to multiple testing. J R Stat Soc Series B Stat Methodol. 1995;57(1):289–300. doi: 10.1111/j.2517-6161.1995.tb02031.x.

22. Gene Ontology Consortium. Gene Ontology Consortium: going forward. Nucleic Acids Res. 2015;43(Database issue):D1049–D1056. doi: 10.1093/nar/gku1179. PMID:25428369.

23. Kanehisa M, Goto S. KEGG: Kyoto Encyclopedia of Genes and Genomes. Nucleic Acids Res. 2000;28(1):27–30. doi: 10.1093/nar/28.1.27. PMID:10592173.

24. Yu G, Wang LG, Han Y, He QY. clusterProfiler: an R package for comparing biological themes among gene clusters. OMICS. 2012;16(5):284–287. doi: 10.1089/omi.2011.0118. PMID:22455463.

25. Wilkerson MD, Hayes DN. ConsensusClusterPlus: a class discovery tool with confidence assessments and item tracking. Bioinformatics. 2010;26(12):1572–1573. doi: 10.1093/bioinformatics/btq170. PMID:20427518.

26. Friedman J, Hastie T, Tibshirani R. Regularization paths for generalized linear models via coordinate descent. J Stat Softw. 2010;33(1):1–22. doi: 10.18637/jss.v033.i01. PMID:20808728.

27. Simon N, Friedman J, Hastie T, Tibshirani R. Regularization paths for Cox’s proportional hazards model via coordinate descent. J Stat Softw. 2011;39(5):1–13. doi: 10.18637/jss.v039.i05. PMID:27065756.

28. Blanche P, Dartigues JF, Jacqmin-Gadda H. Estimating and comparing time-dependent areas under receiver operating characteristic curves for censored event times with competing risks. Stat Med. 2013;32(30):5381–5397. doi: 10.1002/sim.5958. PMID:24027076.

29. Subramanian A, Tamayo P, Mootha VK, Mukherjee S, Ebert BL, Gillette MA, et al. Gene set enrichment analysis: a knowledge-based approach for interpreting genome-wide expression profiles. Proc Natl Acad Sci U S A. 2005;102(43):15545–15550. doi: 10.1073/pnas.0506580102. PMID:16199517.

30. Liberzon A, Subramanian A, Pinchback R, Thorvaldsdottir H, Tamayo P, Mesirov JP. Molecular signatures database (MSigDB) 3.0. Bioinformatics. 2011;27(12):1739–1740. doi: 10.1093/bioinformatics/btr260. PMID:21546393.

31. Liberzon A, Birger C, Thorvaldsdottir H, Ghandi M, Mesirov JP, Tamayo P. The Molecular Signatures Database Hallmark Gene Set Collection. Cell Syst. 2015;1(6):417–425. doi: 10.1016/j.cels.2015.12.004. PMID:26771021.

32. Hanzelmann S, Castelo R, Guinney J. GSVA: gene set variation analysis for microarray and RNA-seq data. BMC Bioinformatics. 2013;14:7. doi: 10.1186/1471-2105-14-7. PMID:23323831.

33. Ru B, Wong CN, Tong Y, Zhong JY, Zhong SSW, Wu WC, et al. TISIDB: an integrated repository portal for tumor-immune system interactions. Bioinformatics. 2019;35(20):4200–4202. doi: 10.1093/bioinformatics/btz210. PMID:30903160.

34. Charoentong P, Finotello F, Angelova M, Mayer C, Efremova M, Rieder D, et al. Pan-cancer immunogenomic analyses reveal genotype-immunophenotype relationships and predictors of response to checkpoint blockade. Cell Rep. 2017;18(1):248–262. doi: 10.1016/j.celrep.2016.12.019. PMID:28052254.

35. Ito K, Murphy D. Application of ggplot2 to pharmacometric graphics. CPT Pharmacometrics Syst Pharmacol. 2013;2(10):e79. doi: 10.1038/psp.2013.56. PMID:24132163.

36. Colaprico A, Silva TC, Olsen C, Garofano L, Cava C, Garolini D, et al. TCGAbiolinks: an R/Bioconductor package for integrative analysis of TCGA data. Nucleic Acids Res. 2016;44(8):e71. doi: 10.1093/nar/gkv1507. PMID:26704973.

37. Deng B, Chen X, Xu L, Zheng L, Zhu X, Shi J, et al. Chordin-like 1 is a novel prognostic biomarker and correlative with immune cell infiltration in lung adenocarcinoma. Aging (Albany NY). 2022;14(1):389–409. doi: 10.18632/aging.203814. PMID:35021154.

38. Liu X, Jia Y, Shi C, Kong D, Wu Y, Zhang T, et al. CYP4B1 is a prognostic biomarker and potential therapeutic target in lung adenocarcinoma. PLoS One. 2021;16(2):e0247020. doi: 10.1371/journal.pone.0247020. PMID:33592039.

39. Park S, Cha H, Kim HS, Lee B, Kim S, Kim TM, et al. Transcriptional upregulation of CXCL13 is correlated with a favorable response to immune checkpoint inhibitors in lung adenocarcinoma. Cancer Med. 2023;12(6):7639–7650. doi: 10.1002/cam4.5460. PMID:36453453.

40. Rehbein G, Simm A, Hofmann HS, Silber RE, Bartling B. Molecular regulation of S100P in human lung adenocarcinomas. Int J Mol Med. 2008;22(1):69–77. doi: 10.3892/ijmm.22.1.69. PMID:18575778.

41. Marshall AJ, D. Q, Draves KE, Shikishima Y, HayGlass KT, Clark EA. FDC-SP, a novel secreted protein expressed by follicular dendritic cells. J Immunol. 2002;169(5):2381–2389. doi: 10.4049/jimmunol.169.5.2381. PMID:12193705.

42. Dieu-Nosjean MC, Antoine M, Danel C, Heudes D, Wislez M, Poulot V, et al. Long-term survival for patients with non-small-cell lung cancer with intratumoral lymphoid structures. J Clin Oncol. 2008;26(27):4410–4417. doi: 10.1200/JCO.2007.15.0284. PMID:18802153.

43. Germain C, Gnjatic S, Tamzalit F, Knockaert S, Remark R, Goc J, et al. Presence of B cells in tertiary lymphoid structures is associated with a protective immunity in patients with lung cancer. Am J Respir Crit Care Med. 2014;189(7):832–844. doi: 10.1164/rccm.201309-1611OC. PMID:24484236.

44. Cabrita R, Lauss M, Sanna A, Donia M, Skaarup Larsen M, Mitra S, et al. Tertiary lymphoid structures improve immunotherapy and survival in melanoma. Nature. 2020;577(7791):561–565. doi: 10.1038/s41586-019-1914-8. PMID:31942071.

45. Helmink BA, Reddy SM, Gao J, Zhang S, Basar R, Thakur R, et al. B cells and tertiary lymphoid structures promote immunotherapy response. Nature. 2020;577(7791):549–555. doi: 10.1038/s41586-019-1922-8. PMID:31942075.

46. Petitprez F, de Reynies A, Keung EZ, Chen TW, Sun CM, Calderaro J, et al. B cells are associated with survival and immunotherapy response in sarcoma. Nature. 2020;577(7791):556–560. doi: 10.1038/s41586-019-1906-8. PMID:31942077.

47. Bindea G, Mlecnik B, Tosolini M, Kirilovsky A, Waldner M, Obenauf AC, et al. Spatiotemporal dynamics of intratumoral immune cells reveal the immune landscape in human cancer. Immunity. 2013;39(4):782–795. doi: 10.1016/j.immuni.2013.10.003. PMID:24138885.

48. Newman AM, Liu CL, Green MR, Gentles AJ, Feng W, Xu Y, et al. Robust enumeration of cell subsets from tissue expression profiles. Nat Methods. 2015;12(5):453–457. doi: 10.1038/nmeth.3337. PMID:25822800.

49. Aran D, Hu Z, Butte AJ. xCell: digitally portraying the tissue cellular heterogeneity landscape. Genome Biol. 2017;18(1):220. doi: 10.1186/s13059-017-1349-1. PMID:29141660.

50. Becht E, Giraldo NA, Lacroix L, Buttard B, Elarouci N, Petitprez F, et al. Estimating the population abundance of tissue-infiltrating immune and stromal cell populations using gene expression. Genome Biol. 2016;17(1):218. doi: 10.1186/s13059-016-1070-5. PMID:27765066.

51. Finotello F, Mayer C, Plattner C, Laschober G, Rieder D, Hackl H, et al. Molecular and pharmacological modulators of the tumor immune contexture revealed by deconvolution of RNA-seq data. Genome Med. 2019;11(1):34. doi: 10.1186/s13073-019-0638-6. PMID:31126321.

52. Collins GS, Reitsma JB, Altman DG, Moons KGM. Transparent Reporting of a multivariable prediction model for Individual Prognosis or Diagnosis (TRIPOD): the TRIPOD Statement. Ann Intern Med. 2015;162(1):55–63. doi: 10.7326/M14-0697. PMID:25560714.

53. Wolff RF, Moons KGM, Riley RD, Whiting PF, Westwood M, Collins GS, et al. PROBAST: a tool to assess the risk of bias and applicability of prediction model studies. Ann Intern Med. 2019;170(1):51–58. doi: 10.7326/M18-1376. PMID:30596875.

