## supplementary_figures for "An inflammation-associated five-gene expression signature stratifies survival and immune states in lung adenocarcinoma: an integrative public-cohort analysis"

Supplementary Figures S1-S5

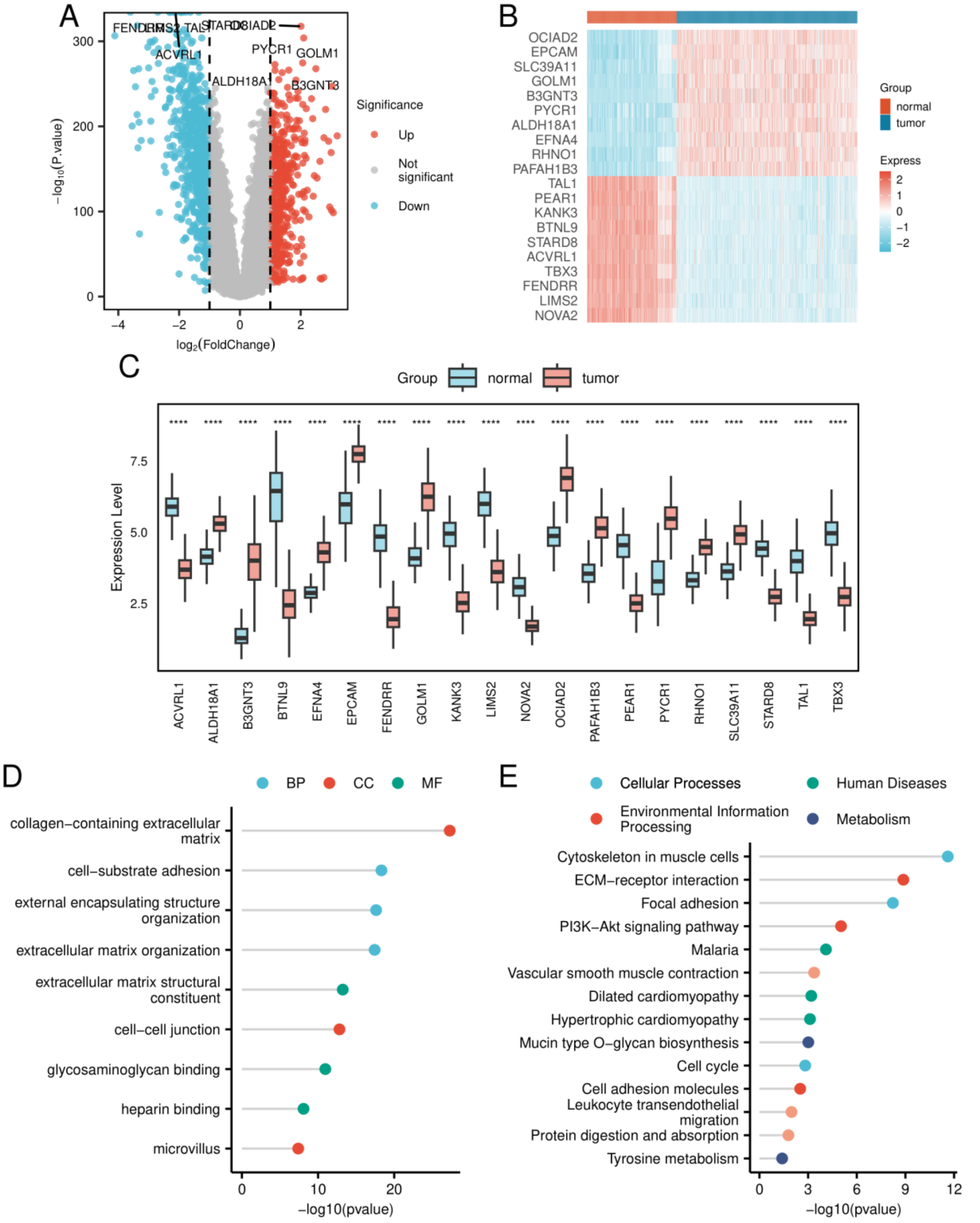


**Supplementary Figure S1.** Identification of differentially expressed genes between LUAD and control lung samples. (A) Volcano plot of LUAD-versus-control differential expression. Red points denote upregulated genes, blue points denote downregulated genes, and gray points denote genes not meeting the significance threshold. (B) Heatmap of selected highly differential genes. (C) Boxplots of 20 displayed genes in LUAD and control samples. (D) GO enrichment of tumor-control differentially expressed genes. (E) KEGG enrichment of tumor-control differentially expressed genes. Differential expression was defined by |log2 fold change|>1 and Benjamini-Hochberg adjusted P<0.05. BP, biological process; CC, cellular component; MF, molecular function; GO, Gene Ontology; KEGG, Kyoto Encyclopedia of Genes and Genomes.

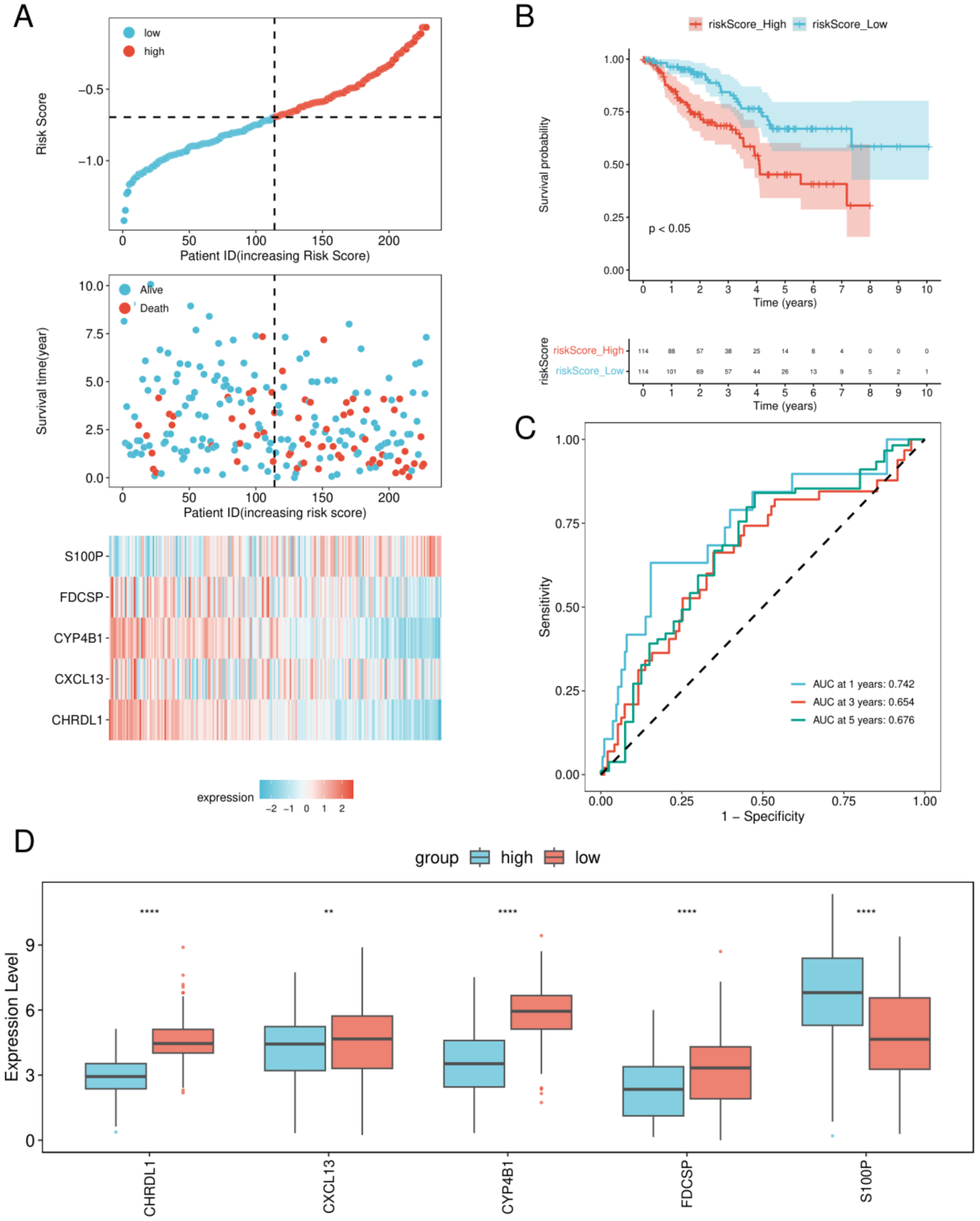


**Supplementary Figure S2.** Internal validation of the five-gene risk score. (A) Risk-score distribution, survival-status distribution, and five-gene expression heatmap in the internal-validation set (n=228). (B) Kaplan-Meier overall-survival curves for high- and low-risk groups. (C) Time-dependent receiver operating characteristic curves at 1, 3, and 5 years; AUCs were 0.7422, 0.6537, and 0.6761, respectively. (D) Expression of the five signature genes in high- and low-risk groups. Significance symbols: ****P<0.0001, ***P<0.001, **P<0.01, and *P<0.05.

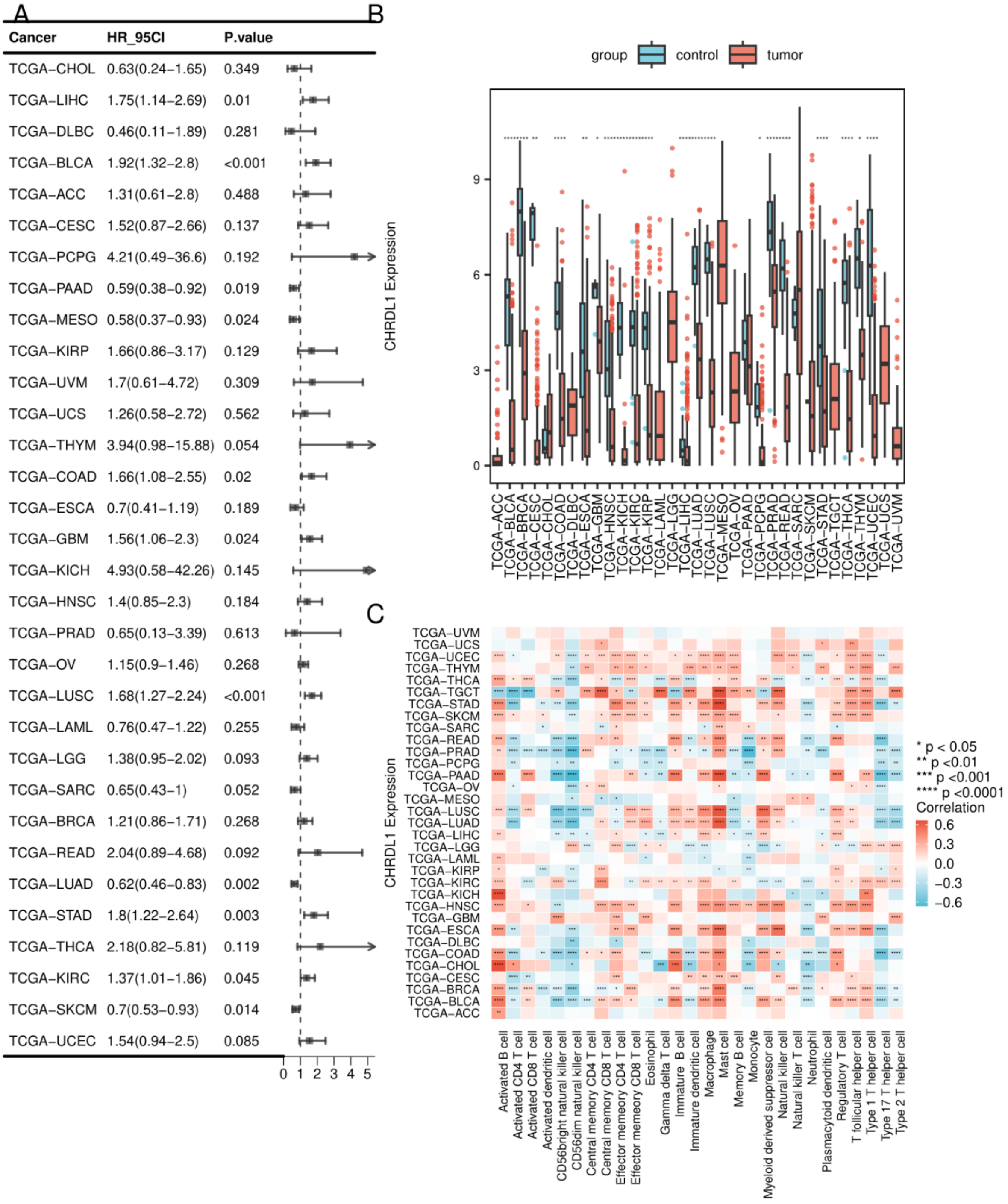


**Supplementary Figure S3.** Pan-cancer analysis of *CHRDL1*. (A) Univariate Cox regression of *CHRDL1* expression and overall survival across TCGA cancer types. (B) Tumor-versus-normal *CHRDL1* expression across cancer types. (C) Correlations between *CHRDL1* expression and 28 immune-cell enrichment scores. In LUAD, the displayed survival estimate was HR 0.62 (95% CI 0.46-0.83; P=0.002).

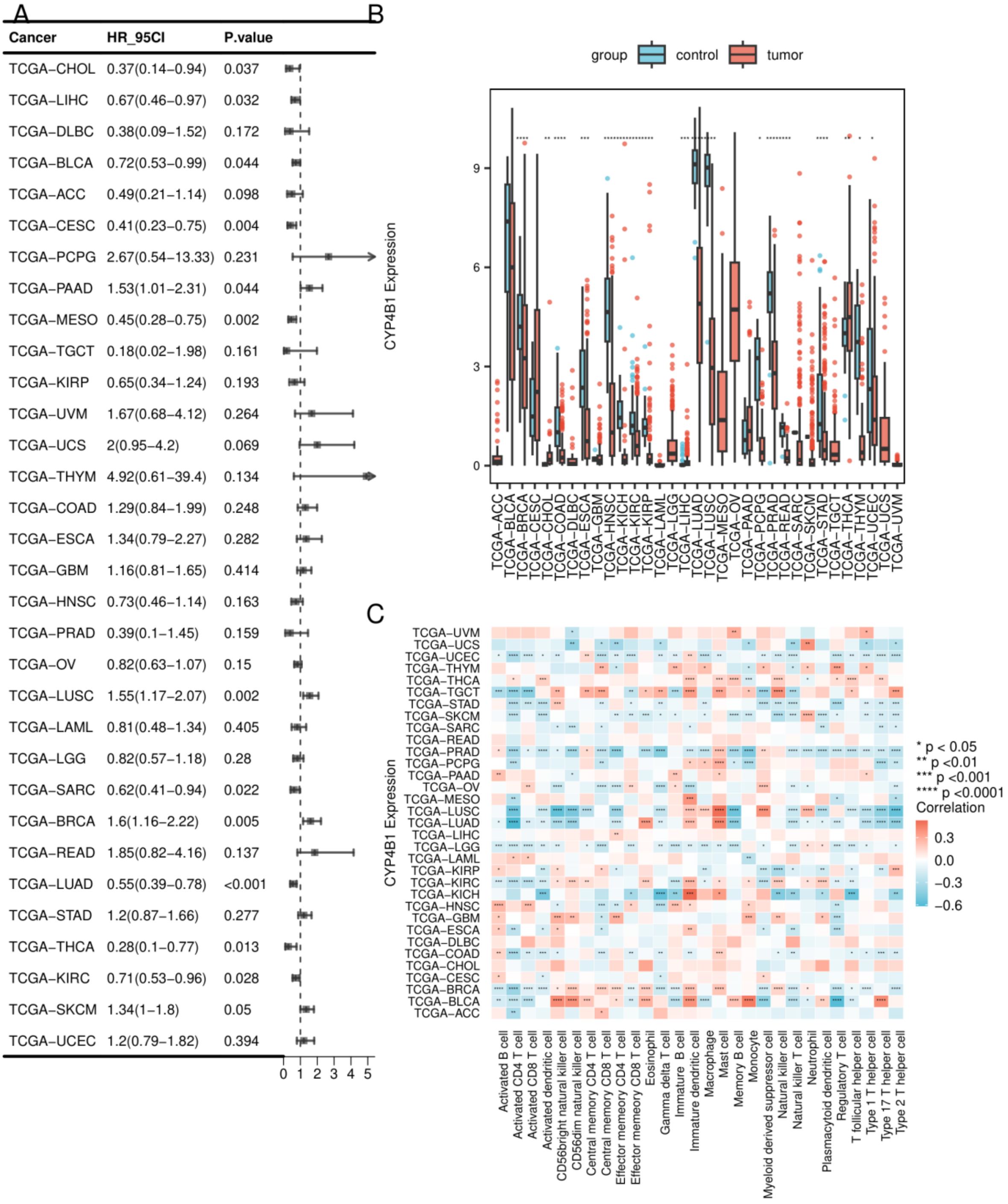


**Supplementary Figure S4.** Pan-cancer analysis of *CYP4B1*. (A) Univariate Cox regression of *CYP4B1* expression and overall survival across TCGA cancer types. (B) Tumor-versus-normal *CYP4B1* expression across cancer types. (C) Correlations between *CYP4B1* expression and 28 immune-cell enrichment scores. In LUAD, the displayed survival estimate was HR 0.55 (95% CI 0.39-0.78; P<0.001).

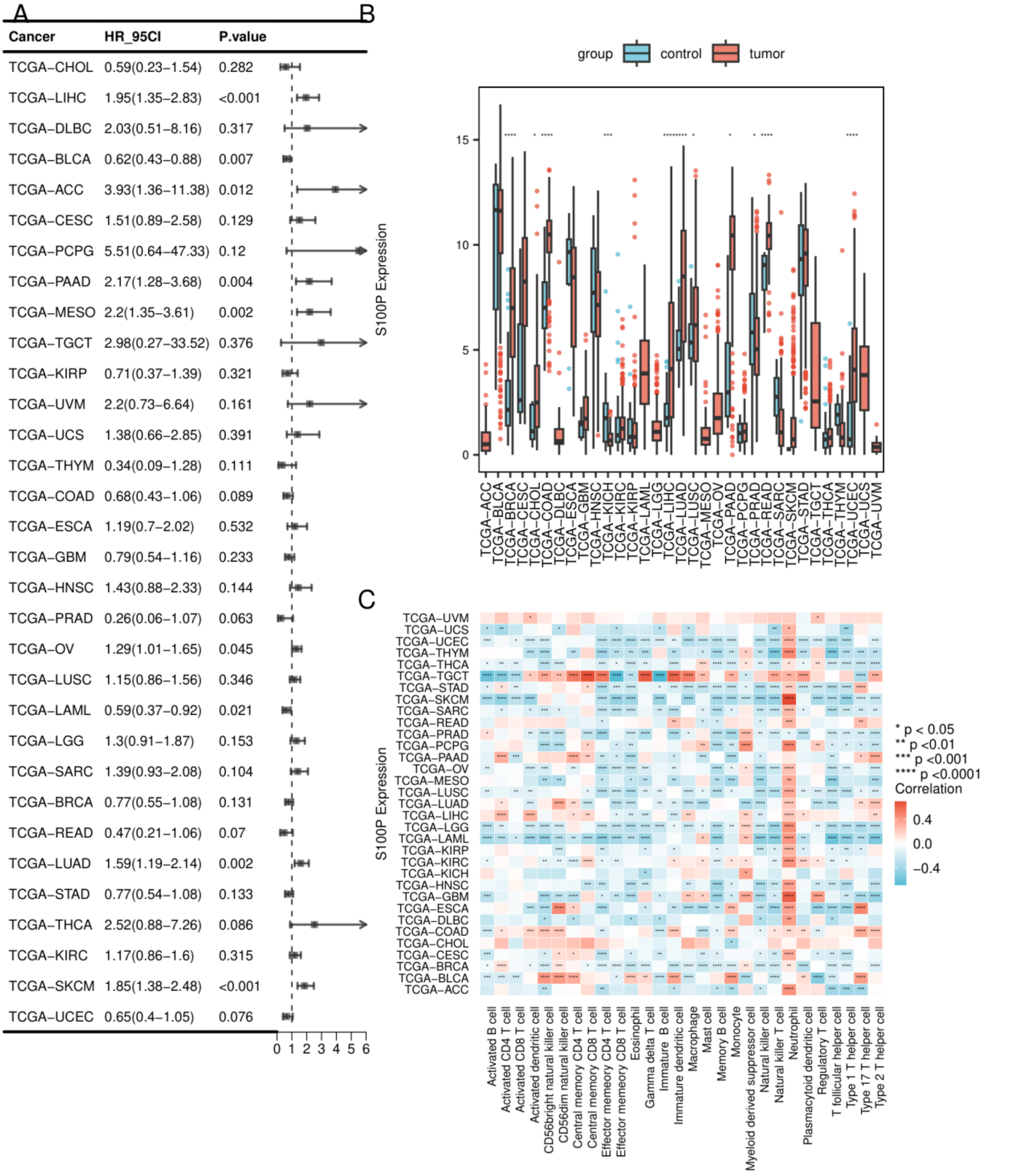


**Supplementary Figure S5.** Pan-cancer analysis of *S100P*. (A) Univariate Cox regression of *S100P* expression and overall survival across TCGA cancer types. (B) Tumor-versus-normal *S100P* expression across cancer types. (C) Correlations between *S100P* expression and 28 immune-cell enrichment scores. In LUAD, the displayed survival estimate was HR 1.59 (95% CI 1.19-2.14; P=0.002).
